# Absent, but not glucocorticoid-modulated, corticotropin-releasing hormone (Crh) regulates anxiety-like behaviors in mice

**DOI:** 10.1101/2024.09.23.614550

**Authors:** Hannah R. Friedman, Lindsey S. Gaston, Li F. Chan, Joseph A. Majzoub

**Author notes:** Address all correspondence to: Joseph A. Majzoub, MD, Division of Endocrinology, Boston Children’s Hospital, 300 Longwood Avenue, Boston, MA 02115. All authors declare they have no conflicts of interest to disclose.

## Abstract

The hypothalamic-pituitary-adrenal (HPA) axis is a well characterized endocrine response system. Hypothalamic Crh in the paraventricular nucleus of the hypothalamus (PVH) initiates HPA axis signaling to cause the release of cortisol (or corticosterone in rodents) from the adrenal gland. PVH-specific deletion of *Crh* reduces anxiety-like behaviors in mice. Here we report that manipulation of PVH *Crh* expression in primary adrenal insufficiency or by dexamethasone (DEX) treatment do not alter mouse anxiety behaviors. In Experiment 1, we compared wildtype (WT) mice to those with primary adrenal insufficiency (*Mrap*KO) or global deletion of Crh (*Crh*KO). We analyzed behaviors using open field (OF) and elevated plus maze (EPM), PVH Crh mRNA expression by spatial transcriptomics, and plasma ACTH and corticosterone after a 15-minute restraint test with ELISAs. EPM analysis showed *Crh*KO mice were less anxious than WT and *Mrap*KO mice, and *Mrap*KO mice had no distinguishing behavioral phenotype. In Experiment 2, we evaluated HPA axis habituation to chronically elevated *Crh* expression by comparing mice treated with 5-8 weeks of DEX with those similarly treated followed by DEX withdrawal for 1 week. All mice regardless of genotype and treatment showed no significant behavioral differences. Our findings suggest that reduced anxiety associated with low *Crh* expression requires extreme deficiency, perhaps outside of those PVH Crh neurons negatively regulated by glucocorticoids. If these findings extend to humans, they suggest that increases in Crh expression with primary adrenal insufficiency, or decreases with exogenous glucocorticoid therapy, may not alter anxiety behaviors via modulation of Crh expression.

## Introduction

The World Health Organization estimates that approximately 300 million people have some form of an anxiety disorder, with this number increasing by 26% during the COVID-19 pandemic.^1^ In general, patients living with anxiety disorders have excessive worry and/or fear. Typically, patients with anxiety disorders have significant disturbances in their thinking which often result in dysregulation of their emotions and/or behavior that may consequently impair daily functions.^1^ There is a wide spectrum in the severity of symptoms patients may experience, and many factors may contribute to the source of their anxiety.

Stressors are defined as conditions in which there is a threat to homeostasis. This includes perceived or anticipated threats established from prior experience and perceived threats within a current situation.^2^ When the body is subjected to these stressors, the stress response incudes autonomic, endocrine, and behavioral components. In the presence of an acute stressor, the autonomic nervous system utilizes both the sympatho-adrenomedullary limb, which secretes epinephrine from the adrenal medulla into the systemic circulation, and the sympatho-neuronal limb, which secretes norepinephrine from sympathetic neurons distributed throughout the body’s organs, to elicit a rapid-onset ‘fight-or-flight’ response.^2–4^ These catecholamines work to restore homeostasis by increasing heart rate, blood pressure, plasma glucose, and oxygen consumption within seconds.^2,5^

Endocrine responses to stressors have a longer time course and are mediated by the hypothalamus, anterior pituitary gland, and adrenal gland, which together constitute the hypothalamic-pituitary-adrenal (HPA) axis.^2–7^ In response to a stressor, corticotropin-releasing hormone (CRH) is released from parvocellular neurons within the medial paraventricular nucleus of the hypothalamus (PVH). Acting through its G-protein coupled CRH receptor-1 (CRHR1), located in corticotrophs of the anterior pituitary gland, CRH stimulates these cells to increase the synthesis and secretion of adrenocorticotropin hormone (ACTH). ACTH stimulates zona fasciculata cells of the adrenal cortex through its G protein-coupled melanocortin-2 receptor (MC2R) to stimulate the synthesis and release of cortisol (or corticosterone in rodents). In addition to their effects to maintain vascular, energy and metabolic homeostasis by promoting vasoconstriction, hepatic gluconeogenesis, proteolysis in muscle, and lipolysis in adipocytes, glucocorticoids cause negative feedback inhibition on CRH in the PVH and ACTH in anterior pituitary corticotrophs via glucocorticoid receptors (GR) to restore the axis to its baseline state.^2,4,6^

The behavioral stress-response has yet to be fully characterized. While CRH plays a role in mouse anxiogenesis, the biological conditions and mechanisms remain unclear.^8^ ^3,9–14^ This study aims to expand upon our and others’ prior work by characterizing the relationship between the body’s endocrine and behavioral stress responses. We have previously shown that deletion of hypothalamic Crh using a Cre recombinase^15^ driven by the expression of transgenic *Sim1* (a transcription factor that regulates PVH development^16,17^), results in both suppression of the HPA stress response and an anxiolytic behavioral phenotype in mice.^18^ This phenotype persisted in mice with *Sim1* deletion of Crh even when they were administered corticosterone-supplemented drinking water, suggesting that the reduction in anxiety was due to loss of hypothalamic Crh rather than the reduction in blood corticosterone concentration.^18^ Another group demonstrated that PVH Crh neurons coordinate rodent behavior in response to stress, as their metaplastic^19^ capability to anticipate^20^ and interpret environmental cues^21^ promotes adaptable behavioral decision making.^20,22^ Through photoactivation/inactivation of PVH Crh neurons in mice, this group found a relationship between PVH Crh activity and the onset of anxiety-like behaviors.^23^ They found that optogenetic stimulation of PVH Crh neurons caused a rapid onset of intense grooming behavior that rapidly ceased with termination of photostimulation^21^. These studies demonstrate that modulation of Crh in PVH neurons controls not only the endocrine response but also aspects of the behavioral response to stressors. This could provide an explanation for the tight coupling between these two aspects of the stress response, which are invariably linked following exposure to a stressor. ^21^

Our present study examines the impact of more subtle, physiological changes in PVH Crh on behavioral responses to stressors by modulating PVH Crh via the negative feedback of glucocorticoids upon PVH Crh expression and comparing these responses to those in *Crh*KO mice. We have used *Mrap*KO mice as a model of high endogenous Crh. These animals have a genetic form of severe, primary adrenal insufficiency due to deletion of *Mrap*, a single-pass transmembrane accessory protein required for signaling of ACTH via Mc2r, causing markedly elevated PVH *Crh* expression.^24^ We have used mice treated with dexamethasone (DEX) as a model of low endogenous Crh. DEX is a potent, synthetic glucocorticoid that suppresses HPA axis activity, as it inhibits both PVH Crh and anterior pituitary ACTH secretion through GR mediation.^25^ ^26^ We administered a supraphysiological amount of DEX in mouse drinking water to WT, *Crh*KO and *Mrap*KO mice, and examined the endocrine and behavioral stress responses in these mice before, during, and after withdrawal from DEX treatment. Surprisingly, we found that while *Crh*KO mice with global deletion of *Crh* have markedly decreased anxiety-like behaviors, modulation of endogenous Crh in mice via alteration in glucocorticoid negative feedback has no effect on behavioral stress responses.

## Methods

### Animal Husbandry

Adult male mice ages 12-16 weeks were used for all experiments. Adrenal function ^27^ including the effect of Crh on adrenal activity ^28^ is sexually dimorphic in mice, such that future studies are warranted in female mice. All mice were bred on a C57BL/6J background and housed in Boston Children’s Hospital’s animal facility following the regulations of the Institutional Animal Care and Use Committee, which approved all experiments. Mice were kept on a 12-hour light/dark cycle (lights on 7 am) in a temperature and humidity-controlled room (21°C with relative humidity 35-64% set point +/- 5%). Animals were kept on a standard chow diet *ad libitium* (<u>Prolab IsoPro RMH3000</u>). *Mrap*KO mice have global deletion of *Mrap* ^24^ and were bred from an *Mrap*KO father paired with a heterozygous (glucocorticoid-sufficient) *Mrap*KO mother.

*Crh*KO mice were generated as previously described from *Crh*KO x *Crh*KO pairings. ^18^

To assure viability of postnatal mice, when either *Crh*KO or heterozygous *Mrap*KO mice were visibly pregnant (around E10.5), mothers were switched from standard drinking water to corticosterone-supplemented drinking water (see below). Mothers and pups were kept on corticosterone-supplemented water until pups were weaned (P21). Mice treated with DEX (see below) were immediately weaned onto DEX-supplemented water (3.6 mcg/mL) and maintained on this treatment for 5-8 weeks.

### Corticosterone Treatment

A 15 mg/mL stock solution was prepared by adding 500mg >92% pure corticosterone (Sigma-Aldrich, C2505-500 mg) to 33 mL pure, non-denatured 200 proof ethanol (Sigma-Aldrich, 493546) and vortexing until the material was completely dissolved. The final stock solution was aliquoted and stored at -20°C until ready for use. Before use, the aliquots were thawed and vortexed until all corticosterone was solubilized and then added into mouse drinking water at a final concentration of 30 mcg/mL, as previously described.^29^

### Dexamethasone Treatment

A 10 mg/mL DEX stock solution was prepared by mixing 100 mg water-soluble DEX (Sigma-Aldrich, D2915-100mg) with 10 mL of tap water. The material was vortexed until completely dissolved. The stock solution was aliquoted and stored at -20°C until ready for use. Before use, the aliquots were thawed and mixed by pipetting up and down and then added to mouse drinking water at a final concentration of 3.6 mcg/mL.

### Behavioral Experiments

#### Open Field

Mice were transferred to Boston Children’s Hospital’s Neurobehavioral Core’s testing facilities (Day 0) and single-housed one week prior to the first testing date to allow the mice to acclimate to their new environment. All experiments were performed starting at 9 am. Open field (OF) experiments were performed (Day 7) in a 42 cm x 42 cm square box and scored using Kinder Scientific Motor Monitor II tracking software. Mice were placed at a 6 o’clock position facing towards the wall at the start time. Each run lasted for 15 minutes. Between each run, each OF box was wiped down with disinfecting/deodorizing wipes. Each mouse was returned to their home cage after testing.

#### Elevated Plus Maze

Elevated plus maze (EPM) experiments were performed (Day 8) on a platform raised 74 cm from the floor. Each arm of the platform was 77 cm in total length. Mice were placed at the center opening so that they were facing the open arms at the start time. Each trial ran for 6 minutes, and the data were automatically scored using Ethovision XT software (version 17.0). EPM heatmaps represent the average amount of time spent by location of each mouse. The average time and location data were generated by the same Ethovision XT software. In Fig. 2, the horizontal arms, denoted with “O”, represent the open arms of the platform, and the vertical arms, denoted by “C”, represent the closed arms on the platform. The center area of the EPM is not enclosed by walls, so it is considered an open area of the maze. Between each run, the maze was wiped down with disinfecting/deodorizing wipes. Each mouse was returned to its home cage after testing. For both OF and EPM experiments, a lamp with an 8.5 W (800 Lumens) light bulb was pointed up at the ceiling for low level ambient lighting.

**Figure 1.**
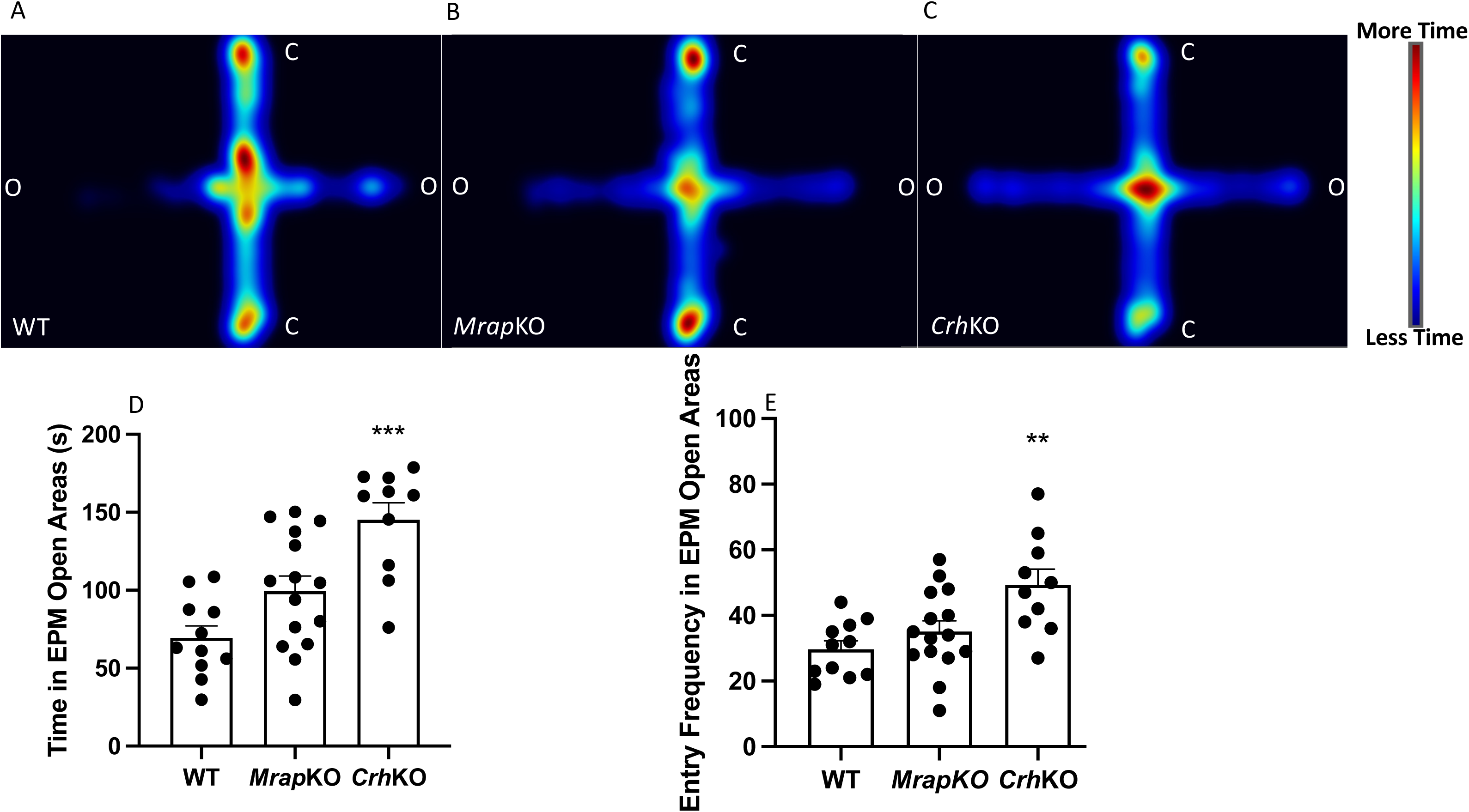
Analysis of anxiety-like behaviors in WT, *Mrap*KO and *Crh*KO mice. (**A-C**) Arial-view heat map of EPM displaying genotype-averaged position x time during 6 min EPM test. In the pictogram, ‘O’ denotes the open arms of the maze while ‘C’ denotes the closed arms. Color gradient indicates the relative time spent within a specified region (Red=maximum, Black = minimum). (**D**) *Crh*KO mice, compared with WT mice, spend the most time in the open areas of the EPM (P<0.0001), whereas *Mrap*KO show no significant difference compared to WT mice (P=0.0595). (**E**) *Crh*KO mice, compared with WT mice, enter the open areas of the EPM more frequently during the 6 min test period (P=0.0015), while *Mrap*KO show no difference compared to WT mice (P=0.5407).

**Figure 2.**
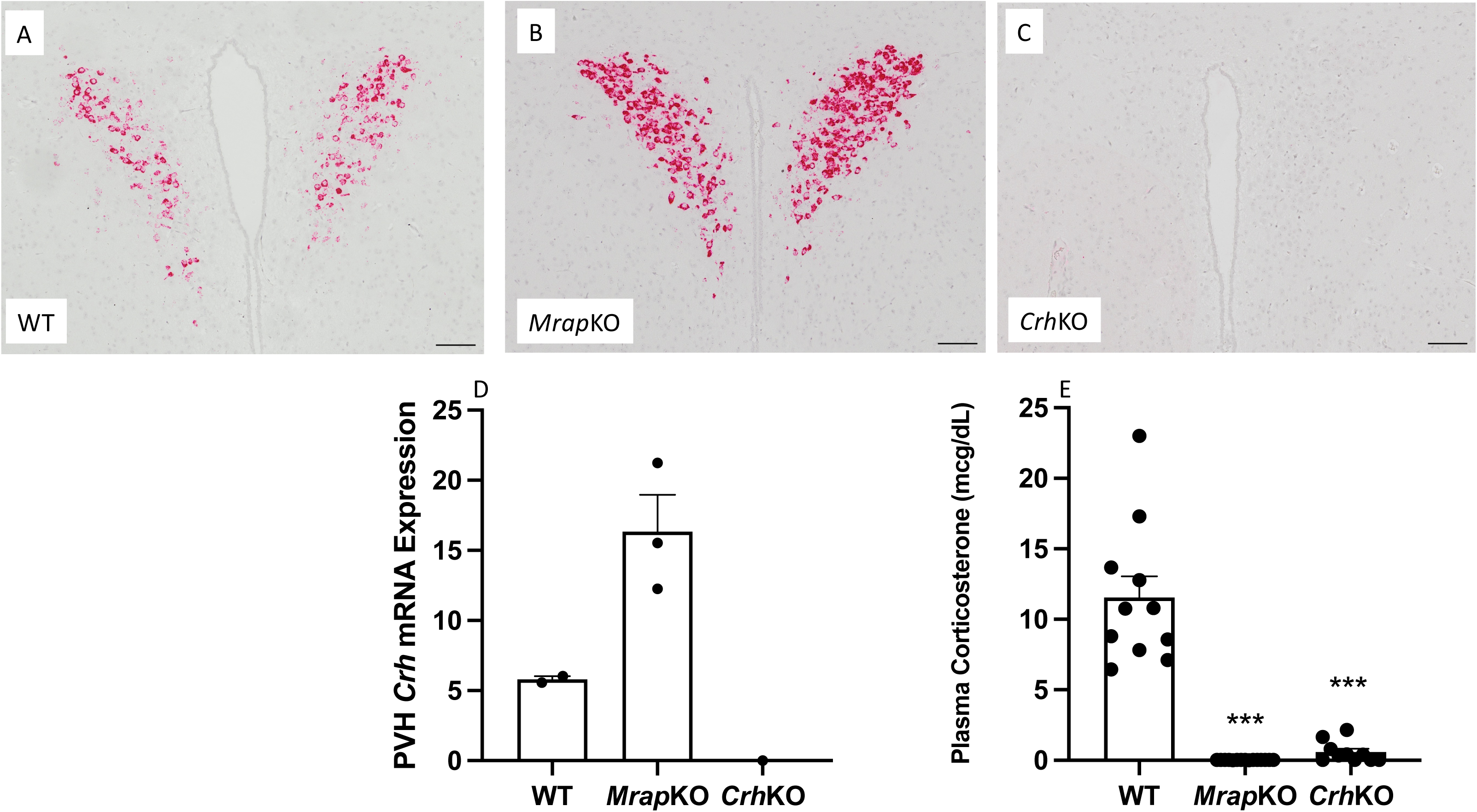
HPA axis status in WT, *Mrap*KO and *Crh*KO mice following 15 minutes exposure to restraint. Representative images of PVH *Crh* mRNA expression in (**A**) WT, (**B**) *Mrap*KO, and (**C**) *Crh*KO mice. Compared to WT, *Mrap*KO have increased, and *Crh*KO have absent *Crh* mRNA expression. (**D**) PVH *Crh* mRNA semi-quantification analysis shows *Mrap*KO mice have a non-significant trend towards increased PVH *Crh* mRNA compared to WT mice (P=0.052). (**E**) Plasma corticosterone concentrations in WT, *Mrap*KO, and *Crh*KO mice immediately after 15 min of restraint stress. Asterisks denote P≤ 0.0001 compared to WT using One-way ANOVA with Bonferroni correction. Compared to WT mice, *Mrap*KO have undetectable, and *Crh*KO have minimal response to restraint stress. Scale bars = 100 μM.

#### Restraint Test and retro-orbital phlebotomy

On Day 9 of Experiments 1 and 2, one day after completing the EPM experiment, mice were restrained in 50cc tubes that previously had the end of the tube cut off for ventilation. After 15 minutes of restraint, mice were subjected to retro-orbital phlebotomy using EDTA-coated capillary tubes and returned to their home cage. Blood samples were immediately placed on ice and then centrifuged at 4°C for 10 minutes at 1500 x g. The plasma was immediately aspirated and stored at -80°C until hormone analysis. A separate group of WT, *Crh*KO heterozygotes, and *Crh*KO homozygotes was restrained for 2 h to verify *Crh* mRNA expression in the *Crh*KO model used in this study that was created by recombineering^18^ and different from our initial *Crh*KO model.^28^

In Experiment 2, due to unavoidable experiment interference, 1 WT untreated animal was restrained for 28 min instead of 15 min. Therefore, data from this animal were removed from both plasma ACTH and corticosterone concentration evaluations. No interference occurred during the prior behavioral experiments, so those data from this animal were retained in OF and EPM analyses.

### Hormone Assays

Plasma was slowly thawed on ice. Corticosterone concentration was measured using a mouse/rat corticosterone ELISA kit (ALPCO; 55-CORMS-E01) as directed by the manufacturer. The lower limit of detection is 0.061 mcg/dL. Animals which had undetectable plasma corticosterone levels were recorded as having a value halfway between 0 mcg/dL and the assay’s lower limit of detection. ACTH levels were measured using ACTH ELISA (ALPCO; 21-ACTH-E01) following the manufacturer’s protocol. The lower limit of detection is 0.22 pg/mL. Additional ACTH 0 calibrator (ALPCO; 11-C-270-10-G25) was used to dilute plasma as necessary. Animals that had undetectable plasma ACTH levels were recorded as having a value halfway between 0 pg/mL and the assay’s lower limit of detection.

### *In situ* Hybridization

Experimental animals were euthanized (Day 10) by rapid decapitation without anesthesia. Brains were removed and fixed in 10% formalin overnight and then transferred to 70% ethanol for histology. Paraffin embedding was performed by either the Histology Core Facility at Beth Israel Deaconess Medical Center or at Boston Children’s Hospital’s Histology Department. Paraffin blocks were sectioned and H&E stained by Boston Children’s Hospital’s Pathology Department. Three 5μM-thick coronal brain sections were sectioned onto each slide, and every other slide was H&E stained.

We surveyed H&E slides under a Nikon E800 microscope to identify adjacent unstained slides that contained PVH. Four unstained slides that were flanked by H&E-identified, PVH-containing brain sections were selected for *Crh* mRNA hybridization using RNAscope (ACDBio). The middle brain section on each of the 4 slides was stained for *Crh* mRNA using a mouse *Crh* probe (ACD #316091) and 2.5 HD RED detection kit (ACD, #316091) following manufacturer’s instructions. Similar brain sections from *Crh*KO mice of all treatment conditions for were analyzed for *Crh* mRNA as a negative control.

For an individual biological sample, the brain section with the highest *Crh* mRNA expression among the 4 stained sections was analyzed densitometrically (see below). These selected slides were then compared across their respective biological replicates. If there was variation in *Crh* expression among biological replicates, the selected slides were ranked from least to greatest *Crh* expression, and the median image was chosen as the final representative image. Semi-quantification of RNAscope images was performed following the RNAscope manufacturer’s protocol for FIJI (Fig. S1, Supplemental Materials^30^). The area of the PVH was determined by first drawing two triangles that were large enough to encompass the *Crh* signal on all slides (Fig. S1A, Supplemental Materials^30^). These same triangles were then transposed onto each image. The areas of the two transposed triangles were measured, and the average area of these triangles was calculated. *Crh* expression was quantified by dividing the area of the *Crh* densitometric signal by the average area of the PVH triangles.

### Genotyping

(see Supplemental Materials^30^

### Statistical Analyses

Statistical analyses of behavioral experiments were performed using Prism GraphPad 10. Means were evaluated using One-way ANOVA multiple comparison testing with the Bonferroni method and compared to untreated WT values. In Experiment 2, Figure 3G used the same WT untreated values from Experiment 1 that were shown in Figure 1D. Asterisks on graphs denote significance (*≤0.05, **≤0.001, and ***≤0.0001). Error bars on graphs represent mean + SEM. Due to multiple undetectable plasma ACTH concentrations in DEX treated animals, Gaussian distribution cannot be assumed. Therefore, one-way ANOVA analysis is unable to test for significance compared to WT untreated animals.

**Figure 3.**
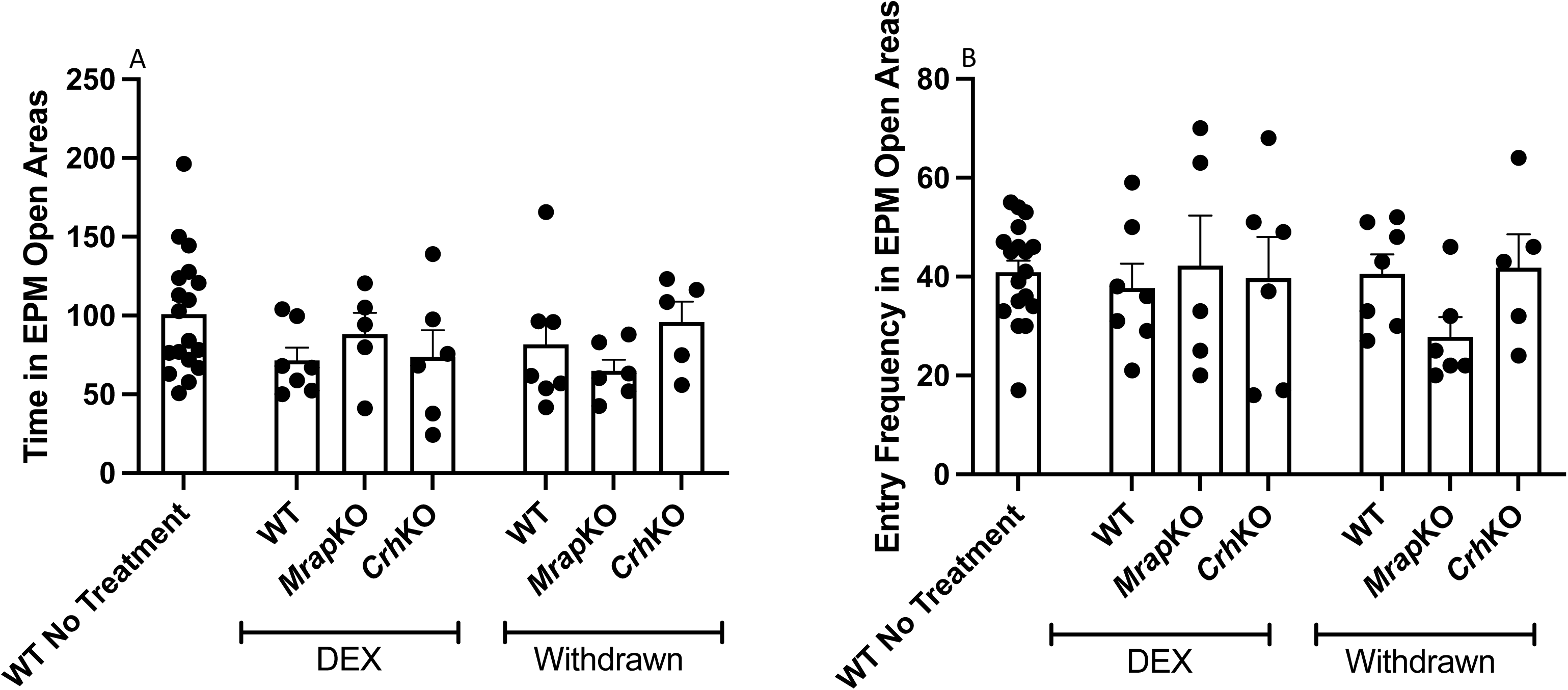
Analysis of anxiety-like behaviors in untreated WT mice compared to WT, *Mrap*KO and *Crh*KO mice either treated with dexamethasone or withdrawn from dexamethasone treatment for 1 week. (**A**) During a 6 min EPM trial, compared to untreated WT mice, all 3 genotypes on dexamethasone treatment or withdrawn from treatment spend comparable amounts of time in the open areas. (**B**) During a 6 min EPM trial, compared to untreated WT mice, all 3 genotypes on dexamethasone treatment or withdrawn from treatment enter the open areas of the maze at the same frequency.

## Results

### *Crh*KO mice lack hypothalamic *Crh* mRNA and have low basal and stress-induced CORT responses

The *Crh*KO model used in this study was created by recombineering^18^, whereas our previously described *CrhKO* model was developed with restriction fragment ligation and retained a *Pgk*neo cassete in the deleted *Crh* locus^28^. We thus first sought to phenotype this novel, recombineered model by assaying hypothalamic *Crh* mRNA expression as well as plasma corticosterone and ACTH levels both in the basal and stressed state in *Crh* WT, *Crh*KO heterozygous, and *Crh*KO homozygous animals. Stress was induced with a 2h physical restraint. Overall, *Crh* mRNA expression significantly differed by both genotype and treatment condition using 2-way ANOVA (Fig S1; p=0.007). *Crh* mRNA increased following 2h restraint in both WT and *Crh*Het animals, and expression was higher in WT vs. *Crh*HET animals under both conditions. As expected, *Crh* mRNA was absent in both basal and stressed *Crh*KO mice (Fig S1, Supplemental Materials^30^).

Corticosterone levels were similarly higher after 2h restraint compared to the basal state in WT and *Crh*HET but not *CrhKO* mice (P<0.0001), though there was no significant effect of genotype nor treatment condition on plasma ACTH levels (P=0.44, Fig S2, Supplemental Materials^30^).

### Influence of endogenous *Crh* and glucocorticoids on behavioral and endocrine responses to stress in WT, *Mrap*KO and *Crh*KO mice. (Experiment 1)

In Experiment 1, we tested whether *Crh* expression links the endocrine and behavioral stress responses. First, we exposed mice to a 15-minute open field (OF) trial followed by a 6-minute elevated plus maze (EPM) trial 24 hours after the OF test. OF total locomotor activity was not different among WT, *Mrap*KO and *Crh*KO mice (WT vs.

*Mrap*KO P>0.999; WT vs. *Crh*KO P>0.999), indicating there were no baseline locomotor abnormalities in the mutant genotypes (Fig. S3A, Supplemental Materials^30^). Further, *Mrap*KO and *Crh*KO mice were not different from WT mice in total distance traveled in the OF center (WT vs. *Mrap*KO P=0.286; WT vs. *Crh*KO P>0.999), time spent in the OF center (WT vs. *Mrap*KO P=0.0719; WT vs. *Crh*KO P=0.836), and number of entries into the OF center (WT vs. *Mrap*KO P=0.462; WT vs. *Crh*KO P>0.999) (Figs. S3B-D, Supplemental Materials^30^). These three measures are indicators of anxiety, suggesting that neither absent (*Crh*KO) nor elevated (*Mrap*KO) *Crh* expression affects these aspects of mouse anxiety-like behavior.

The morning following the OF trial, we exposed mice to a 6-minute EPM trial. Increased time spent and number of entries into the EPM open areas indicate decreased anxiety-like behavior in mice. An arial view heatmap, displaying genotype-averaged position x time during 6-minute trials, showed that *Crh*KO mice (Fig. 1C, D), compared to WT mice (Fig. 1A, D), spent more time in the open areas of the maze (*Crh*KO 145.2±10.8 s vs. WT 69.5±7.6 s; P<0.0001). *Crh*KO mice also entered the open areas more frequently than WT mice during the 6-minute test (*Crh*KO 49.4±4.7 entries vs. WT 29.7±2.5 entries; P=0.0015) (Fig. 1E). Interestingly, *Mrap*KO mice, with elevated PVH *Crh* expression, scored comparably to WT mice in time spent in open areas (*Mrap*KO 99.4±9.6 s vs. WT 69.5±7.6 s; P=0.0595), with a nonsignificant trend towards more time in these areas (Fig. 1B, D). Further, the number of entries into the open areas was not different between *Mrap*KO and WT mice (*Mrap*KO 35.1±3.2 entries vs. WT 29.7±2.5 entries; P=0.541) (Fig. 1E). An increase in latency to enter the open areas of the EPM is associated with increased anxiety-like behavior. Both *Mrap*KO mice (23.6±10.0 s; P=0.911) and *Crh*KO mice (11.9±4.7 s; P>0.999) scored comparably to WT mice (15.0±6.1 s) (Fig. S3E, Supplemental Materials^30^ in this metric.

24 h after completing the EPM trial in Experiment 1, we characterized the HPA axis status in these same WT, *Mrap*KO and *Crh*KO mice after 15 minutes of restraint stress by measuring PVH *Crh* mRNA expression by RNAscope and plasma corticosterone concentrations. Following this stressor, *Mrap*KO mice tended towards increased expression of PVH *Crh* mRNA compared to WT (WT 5.8±0.2 vs. *Mrap*KO 16.4±2.6 arbitrary units; P=0.052) while no *Crh* transcript was detected in *Crh*KO mice (Figs. 2A-D). After 15 minutes of restraint, WT mice plasma corticosterone was 11.6±1.5 mcg/dL, whereas it was undetectable in *Mrap*KO mice (P<0.0001) (Fig. 2E). As we have previously found^18^, *Crh*KO had low plasma corticosterone concentrations following restraint (0.59±0.24 mcg/dL compared to WT, P<0.0001) (Fig. 2E).

### Dexamethasone reduces PVH *Crh* expression and the HPA axis response to stress, and its subsequent withdrawal restores PVH Crh expression and HPA axis responsiveness, but neither treatment affects anxiety-like behaviors. (Experiment 2)

In Experiment 2, we asked if the lack of increased anxiety-like behaviors in *Mrap*KO mice in Experiment 1 was due to habituation to chronically elevated *Crh* expression. To test this, we disrupted HPA axis signaling by administration of dexamethasone (DEX)-supplemented drinking water. We compared two treatment groups to the WT-untreated animals studied in Experiment 1, (Fig. 2A, D). The first group consisted of WT, *Mrap*KO and *Crh*KO mice treated with DEX for 5-8 weeks. The second treatment group consisted of WT, *Mrap*KO and *Crh*KO mice treated with DEX for 5-8 weeks and then withdrawn from treatment for 1 week.

We measured mouse behavior using the same 15-minute OF and 6-minute EPM tests as in Experiment 1. Unlike in Experiment 1, where untreated *Crh*KO mice compared to WT mice had reduced anxiety-like behaviors in OF and EPM tests, following either DEX treatment or its subsequent withdrawal, *Crh*KO mice behaved no differently than untreated WT mice in time spent in EPM open areas (*Crh*KO-DEX 73.8± 16.9 s vs, WT-untreated 100.8± 9.1 s, P=0.616; *Crh*KO-DEX withdrawn 95.8± 13.0 s vs. WT-untreated 100.8± 9.1 s, P>0.999) (Fig. 3A) or entry frequency in EPM open areas (*Crh*KO-DEX 39.7±8.4 entries vs. WT-untreated 40.9±2.4 entries, P>0.999; *Crh*KO DEX-withdrawn 41.8±6.8 entries vs, WT-untreated 40.9±2.4 entries, P>0.999) (Fig. 3B). Similarly, behavior of *Mrap*KO mice in both treatment groups (DEX-treated, and DEX-withdrawn) was not different from that of untreated WT (Fig. 3). All genotypes in both treatment groups (DEX-treated or DEX-withdrawn) had no difference in OF total locomotor activity compared to WT untreated mice, indicating DEX treatment does not interfere with overall mouse locomotion (Fig. S4A, Supplemental Materials^30^). Distance traveled, time spent, and number of entries in the OF center were also not different among all genotypes and treatment groups compared to WT untreated animals (Figs. S4B-D, Supplemental Materials^30^). EPM analysis of latency to open areas also found no significant differences in animals on DEX treatment, nor in animals who were DEX-withdrawn compared to WT-untreated animals (Fig. S4E, Supplemental Materials^30^).

Twenty-four hours after completion of behavioral testing, we analyzed the impact of DEX treatment and its withdrawal on PVH *Crh* mRNA expression and HPA axis hormones after a 15-minute restraint. Compared with *Crh* mRNA expression in untreated WT mice (5.8±0.2 arbitrary units, Figs. 2A, D), DEX-treated WT (Fig. 4A, G, 1.6±0.1 arbitrary units) and *Mrap*KO (Figs. 4B, G, 1.6±1.4 arbitrary units) were lower. Following 1 week of DEX withdrawal PVH *Crh* was higher in both WT (Fig. 4D, G, 6.6±0.4 arbitrary units) and *Mrap*KO (Figs. 4E, G, 7.7±4.3 arbitrary units) mice. As expected, *Crh*KO mice had undetectable *Crh* mRNA irrespective of DEX treatment status (Figs. 4C, F, G). None of the differences in PVH *Crh* were significant between WT and MrapKO genotypes or DEX treatment status, possibly because of the small number of animals studied (Fig 4G; P=0.4948).

**Figure 4.**
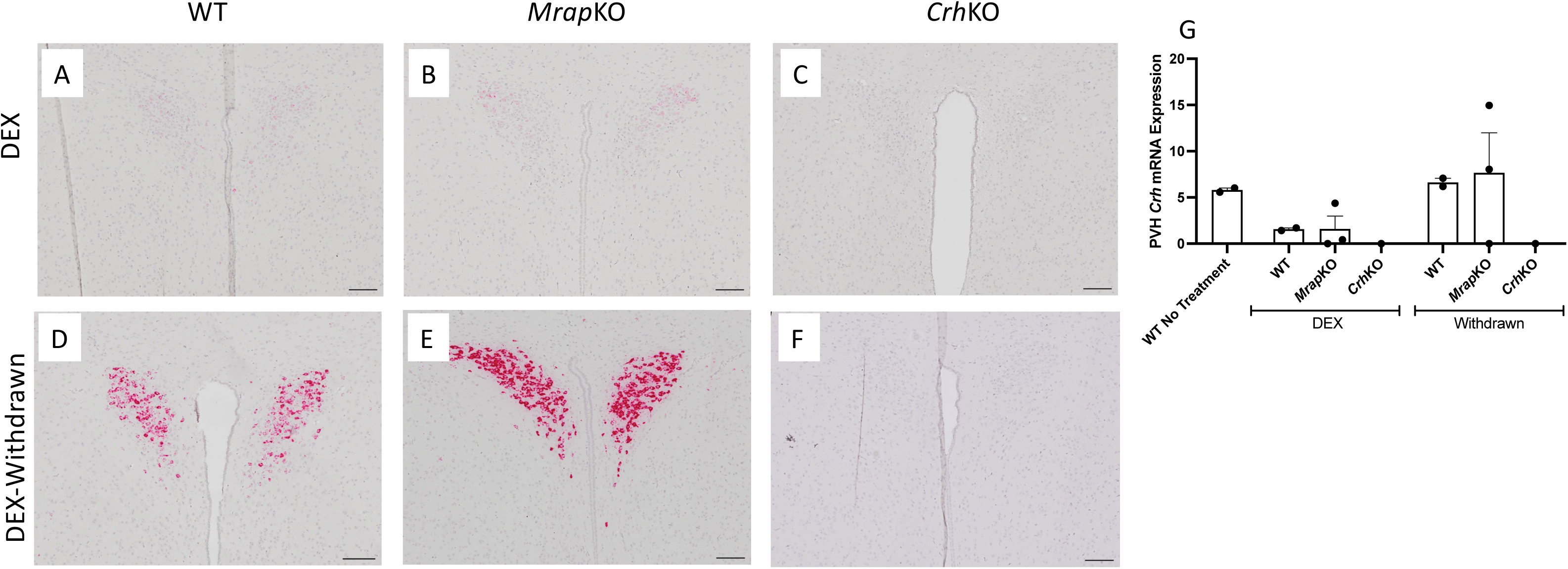
Comparison of paraventricular hypothalamic *Crh* mRNA expression in mice treated with 5-8 weeks of dexamethasone. Representative images of (**A**) WT and (**B**) *Mrap*KO show low *Crh* mRNA expression during dexamethasone treatment. When dexamethasone treatment was discontinued for 1 week, (**D**) WT and (**E**) *Mrap*KO *Crh* mRNA expression increased compared to dexamethasone-treated mice. (**C, F**) *Crh*KO mice have no detectable *Crh* mRNA expression. (**G**) Semi-quantification of PVH *Crh* mRNA shows no significant difference in *Crh* expression when compared to the WT untreated animal values in Figure 2D (P=0.427). Scale bar = 100 μM.

After 15 minutes of restraint stress in WT and *Mrap*KO DEX-treated animals, plasma ACTH was undetectable compared to WT mice not given DEX (410.1±33.9 pg/mL) (Fig. 5A). Plasma ACTH was also very low in DEX-treated *Crh*KO mice (31.8±26.9 pg/mL). One week after DEX withdrawal, WT mice had plasma ACTH concentrations not different from those of WT-untreated mice (WT-withdrawn 833.6±230.6 pg/mL vs. WT-untreated 410.1±33.9 pg/mL; P>0.999) (Fig. 5A), whereas *Mrap*KO mice had significantly elevated plasma ACTH (2399±841.3 pg/mL; P<0.0001) (Fig. 5A). Notably, two DEX-withdrawn *Mrap*KO animals had low (357.9 pcg/mL) or undetectable plasma ACTH after the restraint test. When analyzing the behavior of these two animals, they were not identified as outliers when using the Grubbs’ outlier test (alpha=0.05). Plasma ACTH remained low in DEX-withdrawn *Crh*KO mice (14.6±10.6 pg/mL) (Fig. 5A).

**Figure 5.**
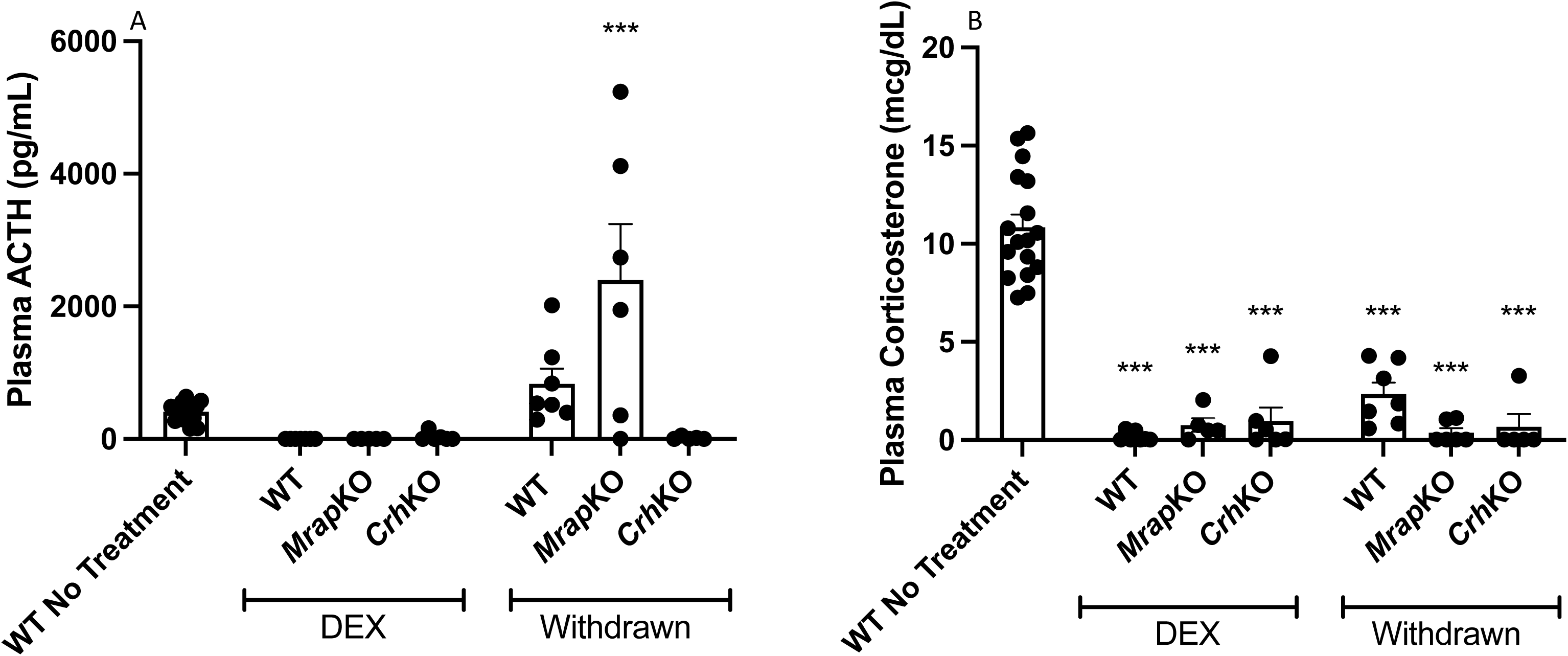
Evaluation of the endocrine response in mice after a 15 min restraint stress test. (**A**) Mice treated with dexamethasone have suppressed plasma ACTH concentrations compared with untreated WT mice. ACTH secretion recovers after 1 week of withdrawing glucocorticoid treatment, with *Mrap*KO mice having significantly elevated plasma ACTH concentrations (P<0.0001). (**B**) Plasma corticosterone concentrations in mice either treated with dexamethasone or withdrawn from dexamethasone. Compared with untreated WT mice, all dexamethasone-treated mice have attenuated corticosterone responses to the restraint stress test, even after withdrawal of dexamethasone 1 week previously. Asterisks denote P<0.0001.

After 15 minutes of restraint stress, plasma corticosterone in WT-untreated mice was 10.9±0.7 mcg/dL (Fig. 5B), similar to that observed in Experiment 1 (Fig. 2E) and was markedly lower in DEX-treated WT mice (0.2±0.1 mcg/dL; P<0.0001) (Fig. 5B). Plasma corticosterone was also very low in DEX-treated *Mrap*KO mice (0.8±0.3 mcg/dL; P<0.0001) and *Crh*KO mice (1.0±0.7 mcg/dL; P<0.0001) (Fig. 4B), with levels similar to those found following restraint without DEX treatment (Fig.1E). Although WT mice withdrawn from DEX treatment had increased plasma ACTH compared to restrained WT mice that had not received DEX (Fig. 4A), corticosterone concentrations after restraint in DEX-withdrawn compared to DEX-untreated restrained WT mice were markedly lower (WT-withdrawn 2.3±0.6 mcg/dL vs. WT-untreated 10.9±0.7 mcg/dL; P<0.0001) (Fig. 4B).

## Discussion

Many studies have explored the role of Crh and its receptor, Crhr1, in anxiety behavior regulation in rodents, ^9,31–35^, including some that have found administration of Crhr1 antagonists results in a reduction of anxiety-like behavioral phenotypes,^10,36^, and another which demonstrated that a Crhr1 antagonist reversed an anxiogenic phenotype in mice with Crh-neuron specific deletion of GABA.^37^ Based on these preclinical data, several pharmaceutical companies developed small molecule CRH receptor 1 antagonists, which unfortunately all failed to reduce anxiety in multiple human clinical trials. ^38^ We previously found that selective deletion of murine *Crh* in the brain regions of *Sim1* expression, including in the PVH, reduces both stressor-induced anxiety-like behaviors and endocrine responses of the HPA axis.^18^ Our findings are consistent with the fact that behavioral and endocrine responses to stressors are usually temporally associated^21^, and suggest that CRH, at least in part, may mediate this linkage. ^23^

In our prior studies of selective deletion of PVH *Crh, Sim1Crh*KO mice were deficient in both hypothalamic Crh as well as in systemic corticosterone, the major mouse glucocorticoid. Thus, their reduced anxiety could have been due to the reduction in either hormone. To evaluate this further we supplemented those mice with short-term systemic glucocorticoid via their drinking water and found that they still exhibited anxiolytic behavior.^18^ We therefore concluded that this behavior was due to hypothalamic Crh, rather than glucocorticoid, deficiency. We undertook the current studies to further examine the impact of manipulation of endogenous Crh via changes in negative feedback by glucocorticoids in mediating stressor-related behaviors. To that end, we have compared wildtype mice to those with global deletion of either *Crh* (*Crh*KO mice) or *Mrap* (*Mrap*KO mice). Mrap is necessary for ACTH function via signaling through the adrenal ACTH/Mc2r receptor. ^24^ Humans with mutations in *MRAP* have a severe form of primary adrenal insufficiency, Familial Glucocorticoid Deficiency Type 2 (FGD2).^39^ Mice with global deletion of *Mrap* have a similar phenotype, with severe glucocorticoid deficiency causing secondary elevation in hypothalamic Crh and pituitary ACTH.^24^ We reasoned that since *Mrap*KO mice have both very low systemic glucocorticoid concentrations and very high hypothalamic PVH *Crh* expression (Fig. 2), they would be useful in understanding whether modulation of one or both of these hormones is involved in stressor-related anxiety behaviors.

In this study, *Crh*KO mice in Experiment 1 exhibited a reduction in anxiety-like behaviors, as they spent more time in the EPM open areas (Fig. 2) and entered the open areas more frequently compared to WT mice (Fig. 2). Their anxiolytic behavioral phenotype was not due to differences in locomotor activity between null mutants and WT mice as determined by both groups’ comparable total distance traveled during the 15-minute OF test (Fig. S3, Supplemental Materials^30^). Thus, the total absence of Crh expression reduces anxiety-like behaviors in mice. We had previously found that a different *Crh*KO mouse line had no anxiolytic phenotype^9^, which was confirmed by another research group using this same mouse. ^40^ However, the mouse model used for those studies^28^ was different from the *Crh*KO mice we used in the present study. The cloned genomic DNA in the original mouse model was obtained from a Balb/c DNA library and resulted in a *Crh*KO mouse on a mixed 129SV/C57BL6 background. The gene-targeting vector replaced the coding region of *Crh* with a neomycin resistance gene that was continuously expressed from a *Pgk* promoter. The *Crh*KO mouse used in the current study was created by breeding a mouse with loxP sites flanking *Crh* exon 2 with an *EIIA*Cre mouse, resulting in germline deletion of the entire *Crh* exon 2.^18^ Further, we have studied this mouse on a pure C57BL6 background instead of on a mixed 129SV/C57BL6 background.

To reverify our new *Crh*KO mouse strain, we used *in situ* transcriptomics to demonstrate our ability to manipulate *Crh* expression in WT and Crh-deficient mice, and affirm our ability to accurately locate and measure PVH *Crh* mRNA. We compared PVH *Crh* expression in WT, *Crh*HET, and *Crh*KO animals in their basal state (Fig. S1, Supplemental Materials^30^) and after a 2 h restraint test (Fig. S2, Supplemental Materials^30^). WT animals had the highest basal PVH *Crh* expression which increased after restraint. *Crh*HET animals had decreased PVH *Crh* expression compared to their respective WT treatment controls and as expected, *Crh*KO animals had no PVH *Crh* expression detected in either. Though WT mice trended towards higher plasma ACTH than *Crh*HET, there was no significant difference among genotypes and treatment groups after 2 h of restraint, possibly because the rise in corticosterone during the 2 h restraint period (Fig. S2, Supplemental Materials^30^) inhibited ACTH production and/or secretion.

The hypothalamus is the central regulator of the body’s endocrine response to an external stressor, as it initiates the HPA axis signaling cascade with Crh. Within the hypothalamus, the highest density of Crh-expressing neurons is located in the PVH,^10^ ^41^ and PVH Crh is activated in the presence of multiple external stressors. In one study, mice were exposed to physical (ether), physiological (high salt), psychological (restraint), immune (lipopolysaccharide), and innate (predator odor) stressors. Out of the 95 brain regions that were examined, the PVH was 1 of 12 areas of Crh expression that had significant increase in Fos expression compared to controls when exposed to all 5 stressors.^41^ The crucial endocrine role of PVH Crh, the dense population of Crh-expressing cells within the PVH, the selective activation of PVH in response to a variety of stressors neurons, and numerous studies that demonstrate the behavioral consequences when PVH Crh is activated or inactivated^18^ ^22^, indicate that PVH Crh has significant importance in rodent stress-response regulation.

Since *Crh*KO mice, with global deletion of Crh, have reduced anxiety-like behaviors, we hypothesized that mice with increased Crh would display an increase in anxiety-like behaviors. However, *Mrap*KO mice, with high PVH Crh expression due to primary adrenal insufficiency (Fig. 2), have normal behavioral responses to stressors (Fig. 1). In fact, analysis of time spent in EPM open areas showed *Mrap*KO mice had a non-significant decrease in anxiety-like behaviors compared to WT mice, spending more time in the open areas. That this might be due to the glucocorticoid deficiency in *Mrap*KO mice is supported by a study that found that WT mice given corticosterone-supplemented drinking water for 4 weeks developed an anxiogenic phenotype.^42^ If an increase in circulating glucocorticoids is an anxiogenic factor, it is reasonable to question if the converse is true. *Mrap*KO mice have undetectable plasma corticosterone which raises the possibility that despite high hypothalamic Crh, their concomitant low glucocorticoid state could result in reduced anxiogenic behaviors, such that the two opposing influences of high Crh and low glucocorticoid in the *Mrap*KO mouse model counterbalance each other, leading to a normal behavioral phenotype.

The amygdala communicates with the hypothalamus to interpret perceived threats.^13,14,37,43,44^ Activation of Crh in central nucleus of the amygdala (CeA) neurons using an AAV-DREADD system increased anxiety-like behavior in mice.^14^ Rats with CeA Crh over expression via stereotaxic viral injection have increased anxiety-like behaviors when evaluated by EPM but not when analyzed with OF. ^43^ Unlike PVH Crh, Crh in the central nucleus of the amygdala (CeA) is stimulated by GC secretion.^44^ The low GC state in *Mrap*KO mice may cause reduced Crh expression in the CeA, leading to a reduction in anxiety that may counterbalance the anxiogenic effect of elevated PVH Crh.

Consistent with our behavioral results in *Mrap*KO mice, another mouse model with high PVH Crh but also high systemic corticosterone did not have a distinct behavioral phenotype.^45^ Those investigators deleted exon 3 of *Nr3c1*, which encodes the glucocorticoid receptor (GR) within the hypothalamus using *Sim1*Cre mice.

Homozygous *Nr3c1*FLOX::*Sim1*Cre+ mice had elevated PVH and CeA Crh expression and elevated systemic corticosterone as a consequence of the PVH failing to appropriately respond to GR-mediated glucocorticoid negative feedback. CeA GR remained intact, as the *Sim1* field does not extend to the CeA, resulting in high systemic corticosterone stimulating CeA Crh.^45^.The absence of increased anxiety-like behaviors in *Mrap*KO (low corticosterone) and in *Nr3c1*FLOX::*Sim1*Cre+ (high corticosterone) mouse models suggests that glucocorticoids may make minimal contributions to the manifestation of anxiety-like behaviors in mice.

It is also possible that the HPA axis in *Mrap*KO mice has somehow undergone physiologic and/or behavioral adaptations to chronically elevated PVH Crh and/or undetectable corticosterone production. A properly functioning stress response system should adapt to learned, recurring internal and external stimuli. ^46^ Some researchers propose neural network habituation as a mechanism for this adaptation, where repeat exposure to a stimulus results in a reduction of stimulus-response pathway transmission.^47^ Anxiety disorders may arise due to perceived threats or disruption in these brain signaling pathways. If a recurring threat or chronic neuronal dysregulation, such as elevated PVH Crh, is then perceived as non-threatening, it could lead to a decrease or cessation of the stress response and therefore any subsequent changes in behavior.^13^ Here we have shown how differing levels of PVH Crh or circulating glucocorticoids do not have a strong relation to anxiety-like behaviors in mice. Though inconclusive, these results together may have broader implications on how investigators should define ‘stress’ in rodents, as many studies use plasma corticosterone concentrations as the sole indicator of the stress response.

In the current study, we tested the hypothesis that the lack of increased anxiety-like behaviors in *Mrap*KO mice was the result of HPA axis adaptations. We studied *Mrap*KO mice that had PVH Crh suppressed by several weeks of DEX treatment, as well as *Mrap*KO mice that had DEX treatment discontinued 1 week prior to behavioral testing. Under these conditions, the endocrine axes of all three genotypes (WT, *Crh*KO, and *Mrap*KO) behaved as anticipated (Figs. 4, 5): *Crh* mRNA expression was suppressed by DEX treatment in WT and *Mrap*KO mice, and recovered 1 week after discontinuation of DEX. Although we did not see total supp ression of PVH *Crh* in mice on DEX treatment, we had previously found that a 70-80% reduction in PVH *Crh* mRNA (by *in situ* hybridization) was associated with an anxiolytic behavioral phenotype in mice.^18^ Restraint-induced plasma ACTH was suppressed in all three genotypes by DEX treatment and recovered above baseline one week after DEX was stopped. Plasma corticosterone was also suppressed during DEX treatment in WT and *Crh*KO and was undetectable in *Mrap*KO, as expected. ^24^ One week after DEX withdrawal, WT corticosterone had not recovered to normal, untreated levels, despite simultaneous ACTH concentrations being above normal.

Both during DEX treatment and 1 week following its withdrawal, none of the three genotypes exhibited any changes in anxiety-like behaviors compared to untreated WT mice (Figs. 3, S4). By testing mice one week after termination of DEX treatment, we had hoped to study them during the acute phase of the increase in PVH Crh. However, it is possible that by the end of the 1-week withdrawal period, these mice had already adapted to their prior, pre-treatment neuroendocrine status, and thus negated our attempt to avoid habituation. It is also possible that global genetic deletion of *Crh* and glucocorticoid-mediated suppression of PVH Crh (and stimulation of CeA Crh) that results in different behavioral outcomes Thus, we have found a reduction in anxiety-like behavior only in mice with genetic manipulation of *Crh* using *Sim1*-Cre in our prior study^18^ and global *Crh* deletion in the present study. In contrast, glucocorticoid-dependent manipulation of Crh expression did not result in any observable behavioral phenotype whether Crh expression was increased or decreased. Our finding that glucocorticoid-induced suppression of PVH Crh is not anxiolytic makes physiological sense—otherwise feedback inhibition of Crh by prior activation of glucocorticoid secretion by a stressor might terminate anxiety behaviors prematurely in response to that stressor. The lack of anxiogenesis we have found in *Mrap*KO mice with elevated PVH Crh requires further exploration.

In our prior studies of anxiolytic behavior in *Sim1*CreCrhKO mice, *Crh* was deleted in ∼70% of neurons within the *Sim1* field. ^18^ Although the strongest activity of *Sim1*Cre is in the PVH, other brain structures that express Crh also express *Sim1* including the anterior/lateral hypothalamic areas and the bed nucleus of the stria terminalis (BNST) ^15^, but not the CeA.^18,45^ Glucocorticoid modulates only a subset of brain neurons that express Crh^48^, so it is possible that glucocorticoid insufficiency in *Mrap*KO mice or excess in DEX-treated mice does not affect all Crh-expressing neurons involved in behavior.

In conclusion, we have found that complete deficiency of Crh is associated with decreased anxiety-like behaviors in mice. These results are consistent with other groups’ work^22^, which establishes Crh as a link between the endocrine and behavioral responses to stress in rodents. The studies which document significant behavioral impact of endogenous Crh, including our own, utilize profound technical manipulations of *Crh* expression (e.g. global deletion, PVH-specific deletion ^18^, optogenetics) that may have forced extreme behavioral outcomes that would otherwise not occur in nature. In our present study, we manipulated *Crh* expression in a way that mimics what could occur in disease (elevation of PVH Crh in primary adrenal insufficiency) or during clinical treatment (PVH Crh suppression during and after glucocorticoid treatment). Although we observed no changes in anxiety-like behaviors following modulation of *Crh* expression by glucocorticoids, it may still be possible that PVH Crh links the endocrine and behavioral responses to stress but in a more complex pathway than we initially hypothesized. PVH Crh along with other limbic regions of the brain that express Crh, may work together to interpret a perceived threat and then produce a behavioral response—explaining why we see a reduction in anxiety when Crh is absent from all brain regions in *Crh*KO mice. Humans with primary adrenal insufficiency^49^ or secondary adrenal insufficiency following exposure to long-term glucocorticoid ^50^ experience increased anxiety. Further characterization of this system could aid in the development of anxiety treatments and improve the quality of life for patients such as these who experience anxiety as a consequence of disorders of their adrenal axis.

## Acknowledgements

We thank Rong Zhang for helpful discussions. Support was provided by NIDDK F32DK131795-02, a Pediatric Endocrine Society Clinical Scholar Award, and a Boston Children’s Hospital Office of Faculty Development/Basic & Clinical Translational Research Executive Committees Faculty Career Development Fellowship (LSG); the Thomas Morgan Rotch Harvard Medical School endowment (JAM); and the Medical Research Council UK and Academy of Medical Sciences fellowship grant (G0802796), a Wellcome Trust grant (217543/Z/19/Z), European Union’s Horizon 2020 research and innovation programme under grant agreement 848077, Barts Charity grant (G-002162) and BBSRC (BB/W018276/1) (LFC). This work was also supported by NICHD IDDRC grant U54 HD090255 at Boston Children’s Hospital.

## Supplemental Materials

Supplemental materials (Figures S1-S4 and genotyping methods^30^) are available at:

## Notes

### Competing Interest Statement

The authors have declared no competing interest.

### Summary of Updates

Funding updated to include NIH funding

