## Supplemental Materials for "Absent, but not glucocorticoid-modulated, corticotropin-releasing hormone (Crh) regulates anxiety-like behaviors in mice" (from another version of this preprint: version 1, 2024-09-24)

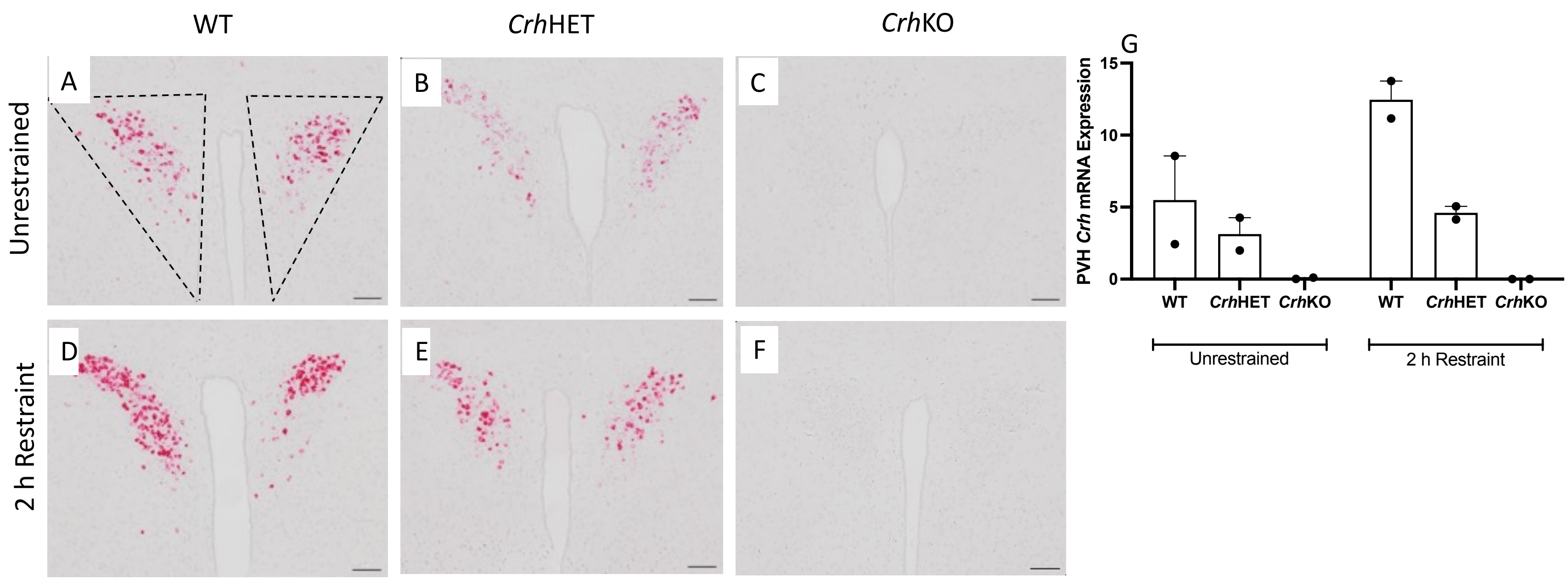


**Figure S1.** Manipulation of PVH Crh expression with different genotypes and 2 h of restraint. (**A**) Representative image of PVH triangle that was transposed onto each stained slide. The area of these triangles was measured and averaged. *Crh* expression was therefore calculated as area of *Crh* signal divided by the average area of the triangles. WT mice have the most PVH *Crh* with basal levels increasing after (**D**) a 2 h restraint test. (**B,E**) *Crh*HET mice have lower PVH *Crh* expression that WT mice. (**C,F**) *Crh*KO mice have undetectable PVH *Crh*. (**G**) Semi-quantitative analysis using One-way ANOVA detects a significant difference of PVH *Crh* expression (P=0.007).

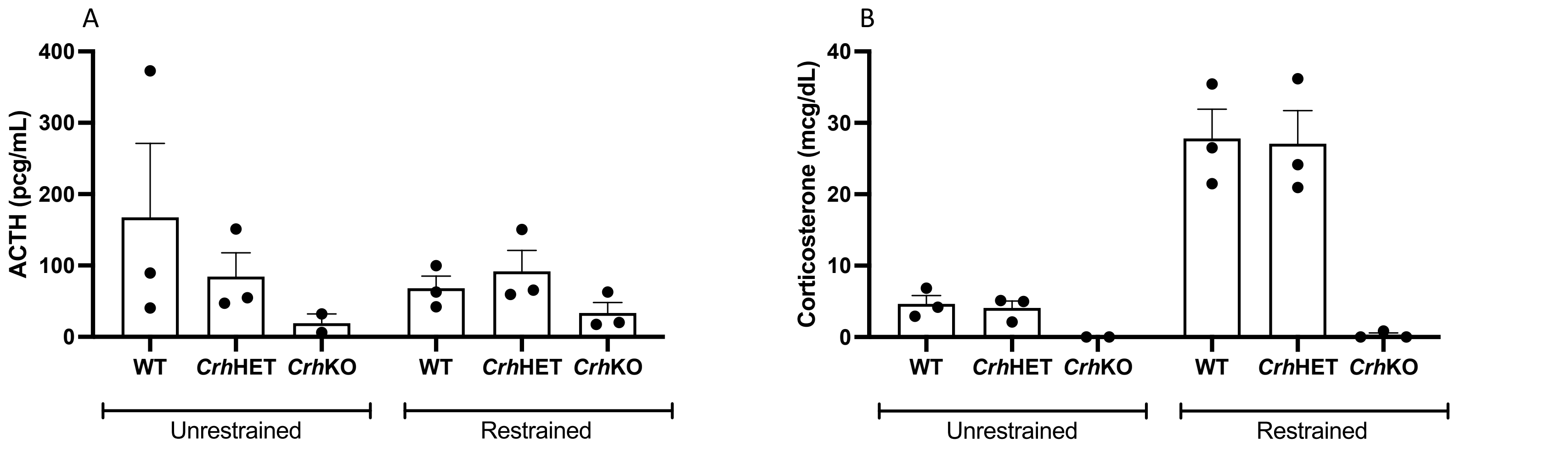


**Figure S2.** Evaluation of the endocrine response in WT, *Crh*HET, and *Crh*KO mice with and without a 2 h restraint test. (**A**) One-way ANOVA analysis among genotypes shows no difference in plasma ACTH concentrations, either unrestrained or after a 2 h restraint test (P=0.4393) (**B**) One-way ANOVA analysis among genotypes shows a significant increase in plasma corticosterone concentrations after a 2 h restraint test (P<0.0001).

Fig. S3

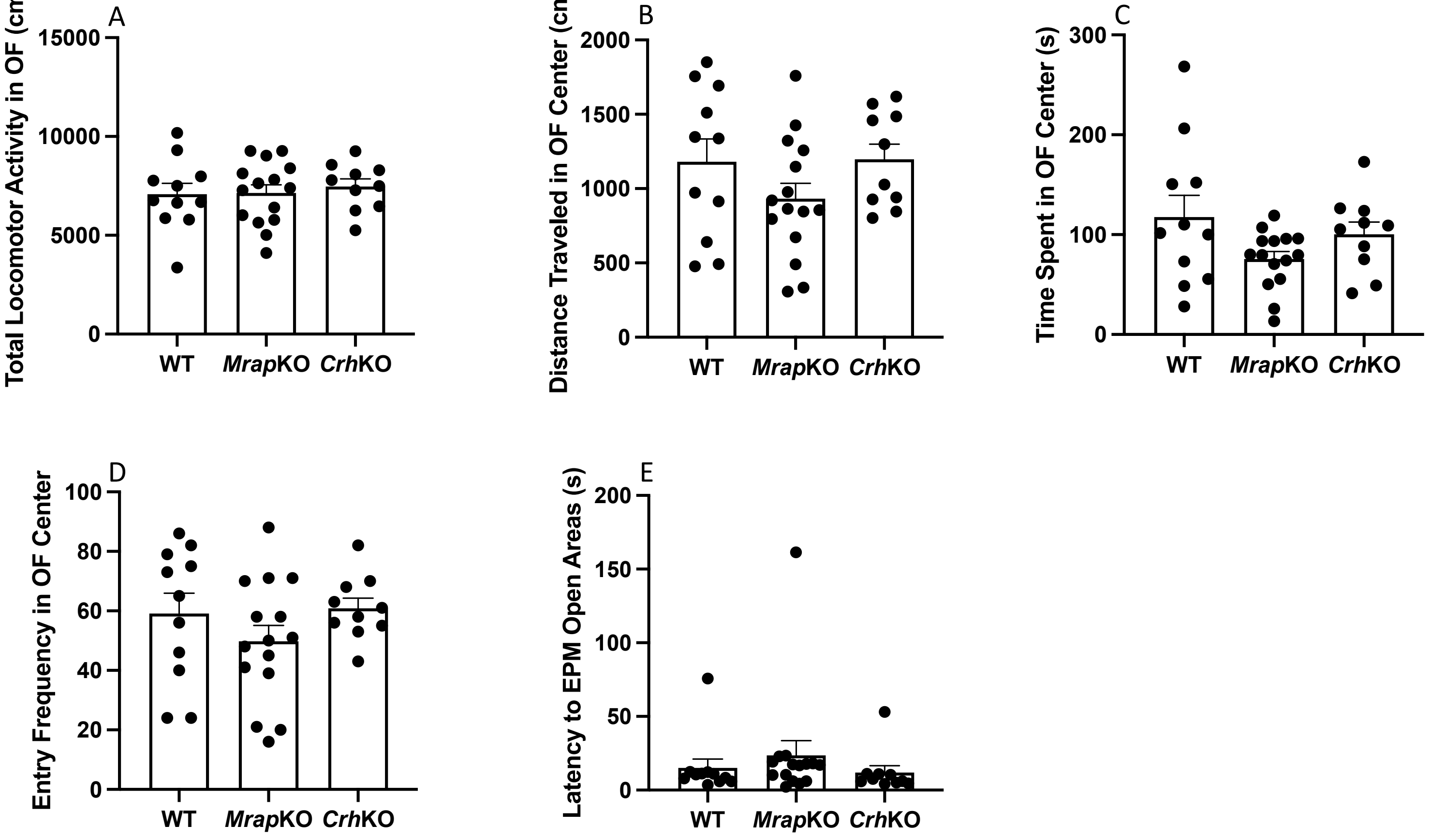


**Figure S3.** Analysis of anxiety-like behaviors in Experiment 1 during a 15-minute open field trial and 6 minute elevated plus maze trial. There are no significant differences among genotypes in the open field trial in (**A**) total distance traveled, (**B**) distance traveled in the center of the open field, (**C**) time spent in the center of the open field, or (**D**) number of entries into the center of the open field. In (**E**) there are no significant differences among genotypes in the latency to enter the open areas of the elevated plus maze.

Fig. S4

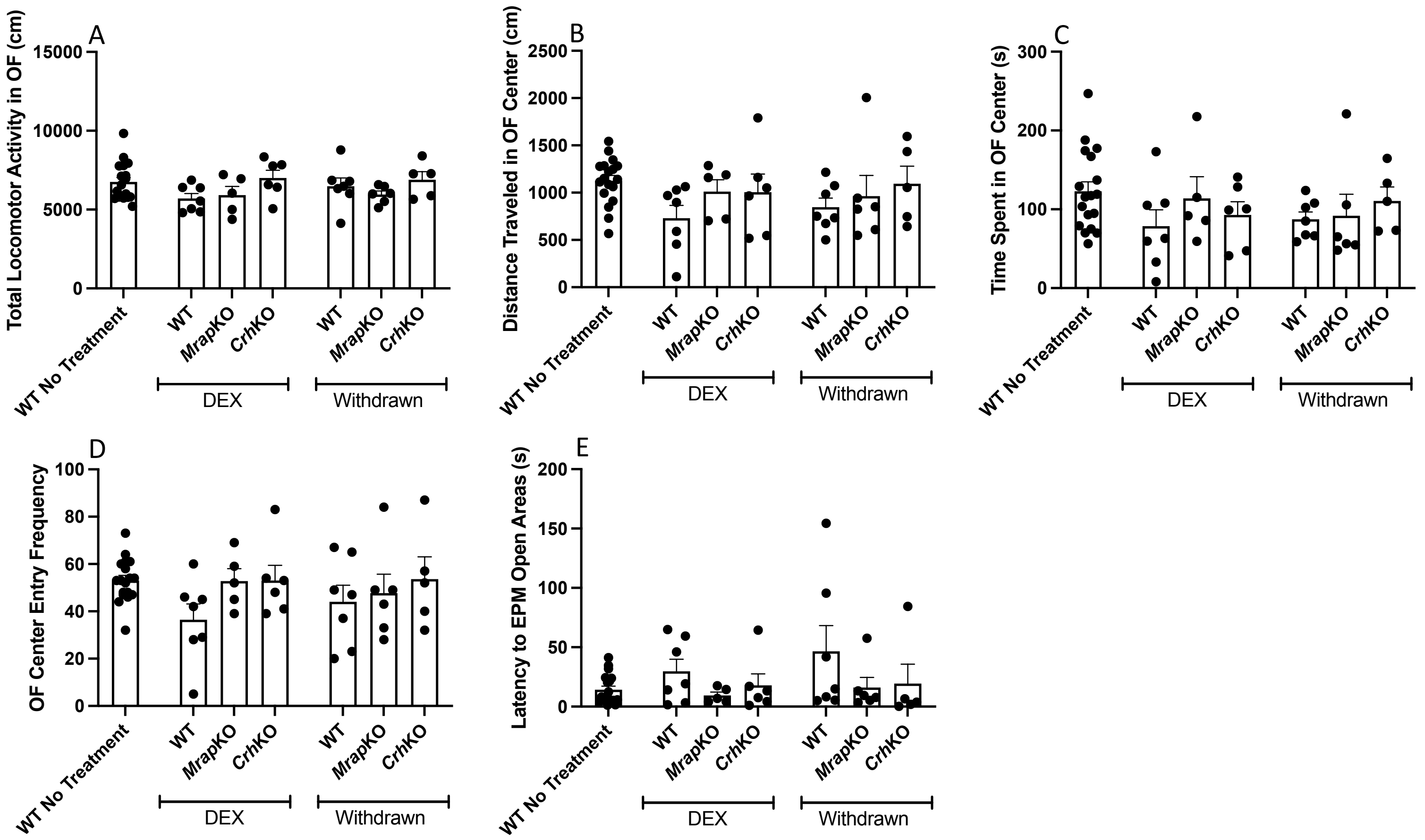


**Figure S4.** Analysis of anxiety-like behaviors in Experiment 2 during a 15 min open field trial and 6 min elevated plus maze trial. Compared to untreated WT mice, among all genotypes, either DEX-treated or DEX-treated and withdrawn 1 week previously, have are no significant differences in the open field trial in (**A**) total distance traveled, (**B**) distance traveled in the center of the open field, (**C**) time spent in the center of the open, or (**D**) number of entries into the center of the open field. In (**E**) there are no significant differences among genotypes or DEX treatment status in the latency to enter the open areas of the elevated plus maze.

Genotyping

Tail DNA was extracted using 0.05 M NaOH, heated at 95°C for 30 min and then neutralized with Tris-HCl (pH 7.4). All genotyping protocols used a 2x Taq RED Master Mix, 1.5 mM $MgCl_2$ (Apex, 5200300-1250) with 10 μM diluted primers (IDT). Primers and PCR protocols for each genotype are as follows:

***Crh*KO**

*Crh* Wild Type Reaction:

*Crh*WT FW: 5'-CAA AGG TTG CTG TGG CTT TAT TTT TCC TCT TCA-3'

*Crh*WT RV: 5'-TTG TCC TCT GAC CTC CAC CCA CTT C-3'

Denature DNA template at 94°C for 2 min. Then cycle 28x at 94°C for 10 s, 66°C for 30 s, and 72°C for 1 min. Finally, extend template for 5 min at 72°C.

*Crh* Knockout Reaction:

*Crh*KO FW: 5'-CCA GCT GCC CAT GTG CTG GA-3'

*Crh*KO RV: 5'-TTG TCC TCT GAC CTC CAC CCA CTT C-3'

Denature DNA template at 94°C for 2 min. Then cycle 28x at 94°C for 10 s, 64°C for 30 s, and 72°C for 1 min. Finally, extend template for 5 min at 72°C.

| | **Expected WT Reaction Band** | **Expected KO Reaction Band** |
|---|---|---|
| ***Crh*WT** | 348bp | No band |
| ***Crh*HET** | 348bp | 375bp |
| ***Crh*KO** | No band | 375bp |

***Mrap*KO**

*Mrap* Wild Type Reaction:

*Mrap*WT F: 5'-GCG TCT CTA GTA GCC TTT GG-3'

*Mrap*WT R: 5'-CTT GTT GGC TTT CAG CTT CC-3'

*Mrap* Knockout Reaction:

*Mrap*KO F: 5'-GAA CTT GCC TGG TCA ACT GTT AAA AGG AC-3'

*Mrap*KO R: 5'-TCT TTA GGC TCA CTT CCT CCA CTG ACC-3'

Both the WT and KO reaction were run as separate reactions with the following same PCR protocol: After initial denaturation of template DNA at 95°C for 2 min, a touch down thermocycling program was used: first cycle at 65°C for 15 s, then decrease by 0.5°C for each subsequent cycle for 10 cycles. Then, extend the template for 15 s at 72°C one time.

Next, cycle at 95°C for 15 s, 62°C for 15 s, and 72°C for 15 s for 25 cycles. Finally, extend at 72°C for 2 min.

| | **Expected WT Reaction Band** | **Expected KO Reaction Band** |
|---|---|---|
| ***Mrap*WT** | 345bp | No band |
| ***Mrap*HET** | 345bp | 450bp |
| ***Mrap*KO** | No band | 450bp |
